# Biomarker discovery in inflammatory bowel diseases using network-based feature selection

**DOI:** 10.1101/662197

**Authors:** Mostafa Abbas, John Matta, Thanh Le, Halima Bensmail, Tayo Obafemi-Ajayi, Vasant Honavar, Yasser EL-Manzalawy

## Abstract

Reliable identification of inflammatory biomarkers from metagenomics data is a promising direction for developing non-invasive, cost-effective, and rapid clinical tests for early diagnosis of IBD. We present an integrative approach to Network-Based Biomarker Discovery (NBBD) which integrates network analyses methods for prioritizing potential biomarkers and machine learning techniques for assessing the discriminative power of the prioritized biomarkers. Using a large dataset of new-onset pediatric IBD metagenomics biopsy samples, we compare the performance of Random Forest (RF) classifiers trained on features selected using a representative set of traditional feature selection methods against NBBD framework, configured using five different tools for inferring networks from metagenomics data, and nine different methods for prioritizing biomarkers as well as a hybrid approach combining best traditional and NBBD based feature selection. We also examine how the performance of the predictive models for IBD diagnosis varies as a function of the size of the data used for biomarker identification. Our results show that (i) NBBD is competitive with some of the state-of-the-art feature selection methods including Random Forest Feature Importance (RFFI) scores; and (ii) NBBD is especially effective in reliably identifying IBD biomarkers when the number of data samples available for biomarker discovery is small.

## Introduction

Inflammatory bowel disease (IBD) refers to disorders that involve chronic inflammation in the gastrointestinal tract. The two main types of IBD are ulcerative colitis (UC), which is characterized by continuous ascending inflammation from the rectum into the colon and periods of relapse and remittance^1^, and Crohn disease (CD), which is characterized by discontinuous skip lesions affecting any part of the gastrointestinal tract^2^. Recent metagenome-wide association studies have implicated some changes in the microbial communities in the gut microbiota with the onset and progression of IBD.^3–6^. However, the precise nature of the changes in the gut microbiota in IBD remains to be fully understood^3^.

IBD, particularly in children, fails to be correctly diagnosed, or diagnosed in a timely fashion, because of the frequency of nonspecific symptoms at the onset of the disease^7, 8.^ Although several non-invasive tests exist for IBD, none has been shown to be capable of diagnosing the two main IBD subtypes with sufficient accuracy^9^. Therefore, a biomarker signature for diagnosing IBD and differentiating between the two major IBD subtypes is highly desirable^8, 10^. Identification of microbial biomarkers is a promising direction, not only for predicting IBD onset but also for predicting IBD risk factors^11^.

Identification of disease microbiomarkers from metagenomics data requires effective computational and statistical methods for determining, from a very large number of candidate biomarkers, a minimal subset of biomarkers that can accurately discriminate between two or more phenotypes (e.g., IBD versus healthy). This task presents several challengs in practice^12^: curse of dimensionality; high degree of sparsity of the metagenomics data; complexity of the underlying biology; limitations of sequencing technology and of methods for determining microbial composition and functional profiles from metagenomic data. To date, several statistical methods have been proposed in the literature to compare an abundance of features (e.g., genes or operational taxonomic units (OTUs)) between two groups^13^. Some of these methods have been designed specifically for RNA-Seq data (e.g., DESeq^14^ and edgeR^15^) while recent tools such as metagenomeSeq^16^ and analysis of composition of microbiomes (ANCOM)^17^ have been developed specifically for metagenomics data, which often exhibits greater sparsity than RNA-Seq data. Machine learning methods for feature selection^18^ offer a promising approach to identifying, from either RNA-Seq or metagenomics data, an optimal subset of the features (potential biomarkers) that can be used to build predictive models that can effectively diagnose a disease or discriminate between disease subtypes.

Recent analysis of microbial ecology networks (MEN) (where the nodes denote microbial taxa and links denote some measure of correlations between the composition of the corresponding taxa) derived from healthy and type 2 diabetes (T2D) groups has shown topological differences between the two networks at the global, module (i.e., sub-networks or communities), and node levels and found that the differences in cluster membership of the nodes in the two networks can serve as biomarkers for T2D^19^. Motivated by these findings, Abbas et al.^20^ hypothesized that MEN corresponding to different phenotypes should exhibit different topologies, and the resulting differences in topology at the node and sub-network levels could be exploited in biomarker discovery. They tested this hypothesis using a framework for network-based biomarker discovery (NBBD).NBBD has two key modules: (i) A network construction module for assembling MEN from the abundance data for microbial taxa (e.g., OTUs); (ii) A node importance scoring module for comparing MEN for the chosen phenotypes and assigning a score to each node based on the degree to which the topological properties of that node differ across two networks. They reported results of experiments with a large dataset of new-onset pediatric IBD metagenomics biopsy samples showing that NBBD could effectively discover IBD biomarkers^20^.

In this study, we build on and extend the results of Abbas et al.^20^ in two aspects (i) We introduce a novel node importance scoring method based on three different node resilience measures^21^ for identifying potential biomarkers. The strength of this approach is that the optimal number of features used to specify a biomarker need not be fixed a priori; (ii) We describe a hybrid approach for integrating network-based and random forest feature importance (RFFI) scores for improving the identification of a minimal subset of features to discriminate between the phenotypes of interest (based on the relative abundance of the microbial taxa represented by the features). We also report results of extensive experiments with several instantiations of the NBBD framework using five different network inference tools, nine node importance scoring functions, and varying number of data samples used to perform feature selection. Our results demonstrate the viability of the NBBD framework for biomarker identification, not only from extremely sparse and high-dimensional data but also from datasets with small number of samples.

## Datasets

BIOM files (see http://biom-format.org) and meta-data (including age, gender, race, disease severity, behavior, and location) for a large cohort IBD dataset^3^ were downloaded from the QIITA (https://qiita.ucsd.edu/) database. The dataset consists of 1359 metagenomics samples including rectal tissue biopsy and fecal samples and each sample has 786 OTUs at the genus level that were extracted using the summarize_taxa.py QIIME script. We filtered the data by discarding fecal samples and samples corresponding to patients with age greater than 18 years. The resulting dataset consists of metagenomic biopsy samples for 657 IBD and 316 healthy control cases, respectively. Thus, each sample (which correspond to a row in the table), is encoded by a tuple of values that represent the relative abundances of the various microbial taxa (indexed by the columns) in the sample. To evaluate our models, we randomly split the data into training and test sets, named DS400 and DS573, such that the training data has 200 healthy and 200 IBD samples, and the test data has 457 healthy and 116 IBD samples. It should be noted that predictive models are often tested on a data distribution that reflects the natural distribution of the different classes. However, in this case, the available IBD and healthy samples do not reflect the natural distribution of IBD and healthy cases in the pediatric population. The prevalence of IBD worldwide has been reported to be close to 0.3% of the population^22^. Hence, given the high degree of class imbalance expected in the natural distribution of data, we anticipate that the reported performance of *all* of the methods in our comparison to substantially overestimate the true performance of the predictive models were they to be deployed in a real-world setting. However, this should not impact the validity of the overall conclusions from our study.

The training data is also used for feature selection (i.e., selecting a subset of features that are most relevant for the classification task). In our experiments, we examined the effect of using a small fraction of the training data for performing feature selection. Specifically, we experimented with the following choices of data for feature selection, which we call the feature selection datasets (FSDS): *DS*50 ⊂ *DS*100 ⊂ *FSD*200 ⊂ *DS*300 ⊂ *DS*400, each with equal numbers of IBD and healthy samples.

### Network-based Biomarker Discovery (NBBD) Framework

We summarize the Network-based Biomarker Discovery (NBBD) framework below: (See Fig. 1, adapted from^20^). Given a feature selection dataset (FSDS) of metagenomics samples in the form of a labeled OTU table: (i) The network construction module, produces two MEN, one from the healty samples, and one from the IBD samples, using the chosen network construction tool (e.g., CoNet^23^); (ii) The node importance scoring module compares the two networks and scores each node in terms of its contribution to the differences between the two networks (as measured using one or more network similarity measures); (iii) The *k* highest scoring nodes provide the *k* features used to train and evaluate binary classifiers for predicting whether or not a given metagenomic sample belongs to a healthy of IBD individual.

**Figure 1.**
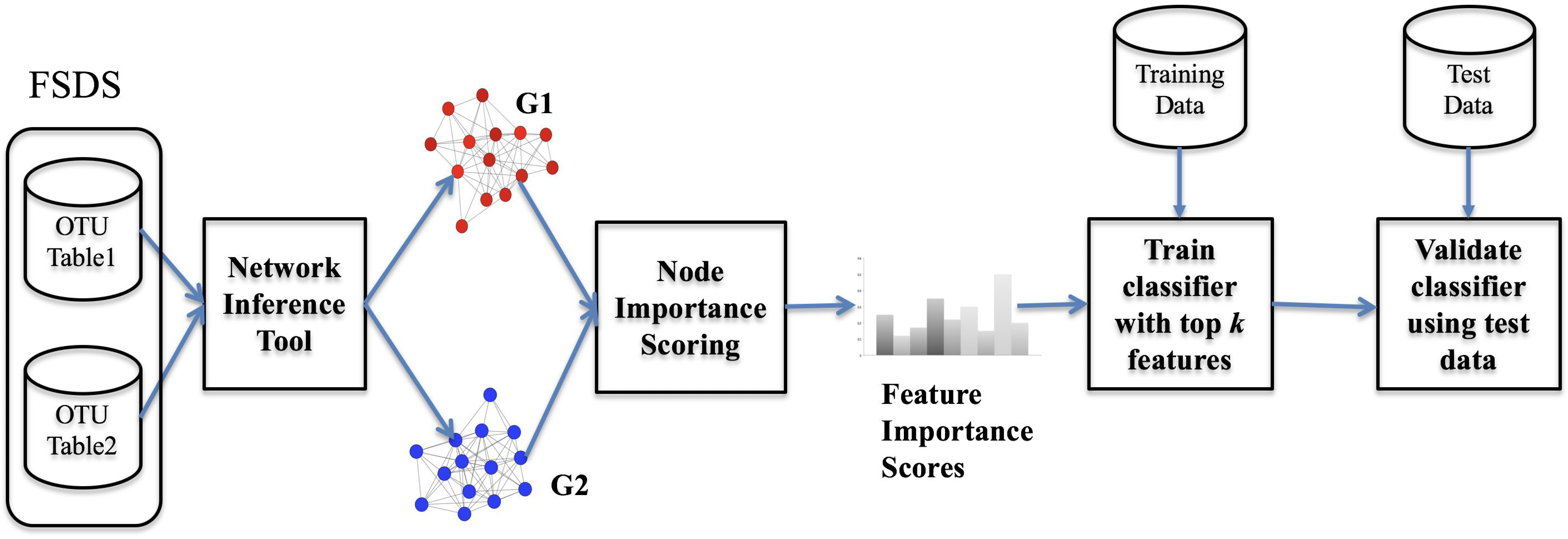
Overview of the NBBD framework. Feature Selection Dataset (FSDS) which is a subset of, or the same as, training dataset in the form of two OTU tables corresponding to two groups of metagenomics samples are first used to construct two networks. The node importance scoring modules compares topological properties of shared nodes in the two networks and outputs scores to prioritize the input features. Top selected features are then used to train and evaluate a classifier.

We evaluated the NBBD framework using five network construction methods and nine node importance scoring methods summarized below.

### Network Construction Methods

We experimented with several widely used methods for constructing MEN from metagenomic data. We used the default parameters of each tool, unless noted otherwise. Each of these methods is briefly described as follows.

- SparCC: Sparse Correlations for Compositional data (SparCC)^24^ infers a network of associations between the microbial species based on the linear Pearson correlations between the log-transformed components (e.g. OTUs). Since log transformation cannot be applied to zeros, which are frequent in microbiome data, zeros are usually substituted with a small value, called pseudo-count. SparCC makes two main underlying assumptions: (i) the number of nodes (e.g. OTUs) is large; and (ii) the underlying network is sparse. We applied the implementation of SparCC included as part of the SPIEC-EASI tool^25^.
- MB: The Meinshausen and Bühlmann (MB) method^26^ is another technique for estimating sparse networks based on estimation of the conditional independence restrictions of each individual node in the graph. The MB method determines the direct neighbors of each target node by finding the smallest subset of nodes such that the target node is conditionally independent of the rest of the networks given the direct neighbors so identified. MB is also implemented in SPIEC-EASI^25^.
- RMT: Random Matrix Theory (RMT) method uses the Pearson correlation coefficient to add an edge between two OTUs if their correlation is higher than a threshold. Instead of using a user-defined threshold, RMT utilizes a procedure based on the Random Matrix Theory to automatically detect a reliable threshold. The method is implemented in the Molecular Ecological Network Analysis Pipeline^27^ available at http://ieg4.rccc.ou.edu/mena. We used the default parameters except for the parameter controlling the number of OTUs that build the network. An OTU was used if it is expressed in at least 25% of the samples. The default value of that parameter is 50% of the samples, and with the parameter set to 50% the method failed to construct the network.
- CoNet: This method infers the association network by combining two complementary approaches to evaluate the significance of the associations^23^. The first approach is an ensemble method of similarity or dissimilarity measures while the second is a novel permutation-renormalization bootstrap method, ReBoot^23^. We followed the procedure described in^28^ to construct the networks for the IBD and healthy phenotypes.
- Proxi: Proxi^29^ is a Python package for proximity graph construction. In proximity graphs, each node is connected by an edge (directed or undirected) to its nearest neighbors according to some distance metric *d*. In our experiments, we set the number of neighbors to seven and used the absolute value of Pearson’s Correlation between two vectors (subtracted from one) as the distance function between two vectors.

### Node Importance Scoring Methods

We considered two approaches for scoring nodes (i.e., features) based on: (i) differences in the topological properties of the nodes in the two networks^20^; (ii) common nodes in the critical attack sets^30^ determined from the two networks. The first approach assumes that a biomarker has different patterns of interactions with other OTUs in healthy and IBD samples. The second approach assumes that biomarkers correspond to a special set of nodes, in the two networks, called a critical attack set^30^ such that the removal of nodes in the critical attack set from a graph results in clustering the network into a number of subnetworks (i.e., microbial communities in the case of MEN).

#### Node Scoring Using Topological Properties

Let *G*_*i*_(*V*_*i*_, *E*_*i*_) and *G*_*j*_(*V*_*j*_, *E*_*j*_) be two graphs constructed using two groups of metagenomics samples (e.g., healthy and IBD). The Node Topological Property Scoring (NTPS) method scores each node *ν* ∈ *V*_*i*_ ∩*V*_*j*_with respect to a node topological property P as follows: *score*^*P*^(*ν*) = |*f*_*P*_(*ν, G*_*i*_) − *f*_*P*_(*ν, G*_*j*_) |, where *f*_*P*_(*ν, G*) is the value of the property P for a node *v* in a graph *G*. In this work, we experimented with the following node properties computed with NetworkX software^31^:

- Betweenness Centrality (btw): Betweenness centrality of a node *v* is defined as 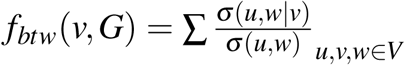, where *σ* (*u, w*) is the total number of shortest paths between *u* and *w* and *σ* (*u, w* |*ν*) is the number of shortest paths between *u* and *w* passing through *ν*.
- Closeness Centrality (cls): Closeness centrality of a node *ν* is given by 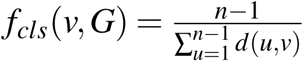, where *d*(*u, v*) is the shortest path distance between *u* and *v* and *n* is the number of nodes that can reach *ν*.
- Average Neighbor Degree (and): The average neighborhood degree of a node *v* is given by 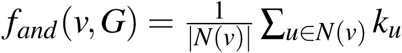, where *N*(*ν*) denotes the set of neighbors of node *ν* and *k*_*u*_ is the degree of node *u* ∈ *N*(*ν*).
- Clustering Coefficient (cc): For unweighted graphs, the clustering coefficient of a node *v* is given by 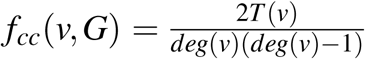, where *T* (*ν*) is the number of triangles that include node *ν* and *deg*(*ν*) is the degree of *ν*.
- Node Clique Number (ncn): The node clique number of a node *ν* is the size of the largest maximal clique containing *ν*. A clique is a subset of nodes such that there is an edge between every pair of distinct nodes.
- Core Number (cn): The core number of a node *ν* is the largest value *k* of a *k*-core containing *ν*, where a *k*-core is a maximal subgraph that contains nodes of degree *k* or more.

#### Critical Attack Set Scoring

Critical Attack Set Scoring (CASS) is based on a node resilience clustering algorithm, NBR-Clust^21^, 30. We briefly describe below, the node resilience measures (specifically the three utilized in this work) before proceeding to describe how they are used to identify biomarkers.

Node-based resilience measures quantify the resilience of a network in terms of the extent of damage (as measured by disruption of connectivity between otherwise connected components or clusters of nodes) caused to the network by the removal of a set of critical nodes (called the attack set)^32^. Because the nodes in the attack set are crucial for maintaining connectivity across the network, removal of the nodes in the attack set can be expected to partition the network into clusters that are isolated from (i.e., disconnected from) each other. Different node resilience measures yield different attack sets with different degrees of sparseness^30^. In this work, we focused on three measures, namely vertex attack tolerance (VAT), integrity, and tenacity.

- The VAT of an undirected, connected graph *G* = (*V, E*) is defined as^32,33^, : 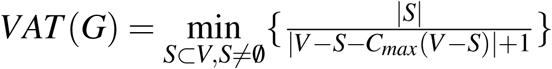, where *S* is an attack set and *C*_*max*_(*V* − *S*) is the largest connected component in *V* − *S*. The goal is to identify small attack sets that consist of nodes that are most cricial in preserving network connectivity.
- Integrity is defined as^34^: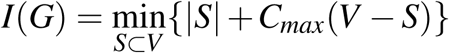. Integrity balances the size of the attack set with the largest connected component in the network resulting from the removal of the attack set. An increase in attack set size can more easily be offset by a decrease in *C*_*max*_, which means that attack set sizes will tend to be larger than with VAT. Generally, the attack set for integrity *S*_*I*_ will include the most crucial nodes (as generated by VAT), plus additional nodes that if removed, make the graph disconnected.
- Tenacity is defined as^35^ :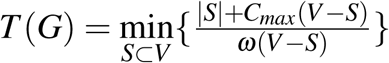, where *ω*(*V* −*S*) is the number of connected components in *V* − *S*. This measure identifies nodes that, if removed, result in partitioning the graph into a large number of components. Thus, the tenacity attack set *S*_*T*_ will include almost all nodes that if removed, can make the graph disconnected.

In order to calculate these resilience measures, we utilized a heuristic known as Greedy betweenness centrality (Greedy-BC)^36^. For a given resilience measure, the Greedy-BC heuristic estimates candidate attack sets by iteratively selecting the node with highest betweenness centrality and removing it from the network. This process results in a node-removal ordering, which is used to calculate all three resilience measures. Each node is then, in order, added to the attack set, with a new graph configuration being generated with each iteration. The resilience measure is updated iteratively after each graph configuration update. The goal is to iteratively optimize the resilience measure. This greedy heuristic can be used to optimize VAT, integrity and tenacity with acceptable accuracy^32^, ^37^. Of the three resilience measures^30^, VAT tends to yield the smallest attack set while tenacity yields the largest. A consequence of using the Greedy-BC heuristic is that the three attack sets are related as follows: *S*_*V*_ ⊆*S*_*I*_ ⊆*S*_*T*_.

To select features for training IBD classifiers, we apply the NBR-Clust algorithm separately to the the IBD and Healthy networks to obtain the critical attack sets for healthy (*G*_*H*_) and IBD (*G*_*D*_) samples. We then select the nodes that are shared by the critical attack sets of both graphs.

### Identification and Evaluation of IBD Biomarkers

Given a training dataset DS400, a feature selection dataset (e.g., DS50), a test set DS573, a feature selection method (FSM), and the number of selected features *k* ∈ {10, 20, 30, 40, 50, 60} : First, we applied the FSM to the feature selection data to determine top *k* features. Then, we generated variants of the training and test data with only the selected features and used them to train and estimate the performance of a Random Forest (RF)^38^ classifier. In each case, the input to the classifier consists of the relative abundance of the microbial taxa represented by the selected features. In our experiments, we used RF classifiers implemented in Scikit-learn^39^ with the number of estimators set to 500 trees.

In addition to our proposed network-based feature selection methods, we considered the following traditional and commonly used feature selection methods: (i) Filter-based feature selection using Information Gain (IG) and F-Statistic (FStat); (ii) Recursive Feature Extraction (RFE) that uses LASSO^40^ estimator for estimating the importance of features and removes the lowest ranked 10 features at each iteration; (iii) RF Feature Importance (RFFI) which is an embedded feature selection method where the FS data are used to train a RF classifier with 500 trees, and feature importance scores are then inferred from the learned model as suggested by Breiman^38^.

We report the predictive performance of all IBD classifiers considered in this study as measured using Accuracy (ACC), Sensitivity (Sn), Specificity (Sp), Matthews Correlation Coefficient (MCC), and Area Under ROC Curve (AUC)^41^.

## Results

### Feature Selection Improves the Predictive Performance of RF Classifiers

Table 1 reports the performance of top (in terms of highest AUC and smallest number of selected features) RF classifiers using five different feature selection datasets as well as using all input features (FSM = None). For RF classifier without feature selection method, the AUC is 0.74. Using the smallest feature selection dataset (DS50), the three traditional feature selection methods yield RF classifiers with better AUC scores. The highest observed AUC corresponds to a RF classifier trained using the top 50 features selected using RFFI method. On the other hand, when using the largest feature selection dataset (DS400), all feature selection methods yield models with AUC better than the baseline model with no feature selection. Interestingly, RFFI seems to benefit substantially by increase in the size of the feature selection dataset since it returns only 20 features that are as discriminative as the 50 features determined using DS50.

**Table 1.** Performance of the top (in terms of highest AUC and smallest number of selected features) performing RF classifiers for different choices of feature selection dataset and traditional feature selection methods.

| FSDS | FSM | # Features | ACC | Sn | Sp | MCC | AUC |
| --- | --- | --- | --- | --- | --- | --- | --- |
| DS50 | None | NA | 0.66 | 0.64 | 0.75 | 0.31 | 0.74 |
|  | IG | 60 | 0.65 | 0.62 | 0.78 | 0.32 | 0.76 |
|  | FStat | 60 | 0.63 | 0.64 | 0.62 | 0.21 | 0.69 |
|  | RFE | 40 | 0.69 | 0.66 | 0.78 | 0.36 | 0.79 |
|  | RFFI | 50 | 0.68 | 0.65 | 0.82 | 0.38 | 0.80 |
| DS100 | None | NA | 0.66 | 0.64 | 0.75 | 0.31 | 0.74 |
|  | IG | 60 | 0.65 | 0.62 | 0.74 | 0.29 | 0.75 |
|  | FStat | 20 | 0.68 | 0.66 | 0.72 | 0.32 | 0.74 |
|  | RFE | 50 | 0.66 | 0.62 | 0.81 | 0.35 | 0.78 |
|  | RFFI | 40 | 0.68 | 0.65 | 0.80 | 0.37 | 0.79 |
| DS200 | None | NA | 0.66 | 0.64 | 0.75 | 0.31 | 0.74 |
|  | IG | 20 | 0.69 | 0.68 | 0.73 | 0.34 | 0.79 |
|  | FStat | 50 | 0.68 | 0.67 | 0.72 | 0.31 | 0.75 |
|  | RFE | 60 | 0.65 | 0.62 | 0.76 | 0.30 | 0.78 |
|  | RFFI | 20 | 0.67 | 0.63 | 0.81 | 0.36 | 0.79 |
| DS300 | None | NA | 0.66 | 0.64 | 0.75 | 0.31 | 0.74 |
|  | IG | 30 | 0.69 | 0.66 | 0.80 | 0.38 | 0.80 |
|  | FStat | 60 | 0.68 | 0.67 | 0.75 | 0.34 | 0.76 |
|  | RFE | 60 | 0.68 | 0.65 | 0.80 | 0.36 | 0.79 |
|  | RFFI | 30 | 0.68 | 0.64 | 0.81 | 0.37 | 0.79 |
| DS400 | None | NA | 0.66 | 0.64 | 0.75 | 0.31 | 0.74 |
|  | IG | 60 | 0.64 | 0.61 | 0.73 | 0.28 | 0.75 |
|  | FStat | 40 | 0.70 | 0.69 | 0.72 | 0.34 | 0.76 |
|  | RFE | 60 | 0.64 | 0.62 | 0.73 | 0.28 | 0.76 |
|  | RFFI | 20 | 0.69 | 0.68 | 0.76 | 0.36 | 0.80 |

We find that some feature selection methods (e.g., IG) are sensitive to changes in the FSDS. For example, the best subset of features returned using the IG filter is with DS300. On the other hand, with DS400 (which includes all instances in DS300), the IG filter fails to determine a good subset of selected features. We suspect that the biomarkers identified using such unstable feature selection methods are likely to be unreliable.

### Performance of Network-based Feature Selection Methods

Results in Table 1 demonstrate the superior performance of RF feature importance for identifying a small subset of discriminative features from metagenomics data which is widely acknowledged in the literature^42,43^. Here, we report results of experiments (using the framework in Fig. 1) designed to address the following questions: (i) which network inference tool learns graphs that could be suitable for our network-based feature selection method?; (ii) how do the results of network-based feature selection using different Node Topological Property Scoring (NTPS) and Critical Attack Set Scoring (CASS) compare to each other as well as to results in Table 1?

First, for each of the five FSDS considered in our experimented and using graphs generated by five NIMs, we evaluated our NBBD framework using six topological properties for NTPS approach and *k* identified biomarkers for *k* ∈ {10, 20, 30, 40, 50, 60}. A total of 900 experiments were conducted and are reported in Tables S1-S5. Table 2 summarizes all these tables for results obtained using DS50 by reporting the performance of top performing (in terms of highest AUC and smallest number of selected features) RF classifiers. Table 2 reveals the following interesting observations: (i) Models using networks generated by CoNet, Proxi, and RMT achieve performance comparable to that of best performing models in Table 1 using RFFI and RFE feature selection; (ii) The AUC of the top performing models obtained using RMT graphs are consistently good (i.e., AUC scores in the range 0.77-0.78), while other NIMs yield top performing models with a wider range of AUC scores; (iii) There is no single topological property that can be used to train RF classifiers that outperform their counterparts trained using other properties. However,the topological properties that work best appear to depend on the network construction method used. For example, CoNet and Proxi based models achieve their highest AUC scores using ‘and’ and ‘cn’ properties, respectively. Even though RMT based models have almost the same AUC for all six different topological properties, the method seems to work best with ‘cc’ property since it reaches the highest AUC score of 0.78 using only 20 features whereas it requires at least 50 features using other properties.

**Table 2.** Performance of top (in terms of highest AUC and smallest number of selected features) performing RF classifiers for combinations of different choices of Network Inference Method (NIM) and network-based feature selection using different properties for Node Topological Property Scoring. All results were obtained using DS50 as the feature selection dataset.

| NIM | FSM | # Features | ACC | Sn | Sp | MCC | AUC |
| --- | --- | --- | --- | --- | --- | --- | --- |
| CoNet | and | 50 | 0.69 | 0.66 | 0.81 | 0.38 | 0.79 |
|  | btw | 30 | 0.67 | 0.64 | 0.78 | 0.34 | 0.78 |
|  | cc | 60 | 0.67 | 0.65 | 0.76 | 0.33 | 0.77 |
|  | cls | 60 | 0.66 | 0.63 | 0.77 | 0.32 | 0.74 |
|  | cn | 60 | 0.67 | 0.64 | 0.79 | 0.35 | 0.78 |
|  | ncn | 60 | 0.66 | 0.64 | 0.73 | 0.31 | 0.75 |
| MB | and | 50 | 0.64 | 0.63 | 0.69 | 0.26 | 0.71 |
|  | btw | 50 | 0.65 | 0.62 | 0.77 | 0.31 | 0.74 |
|  | cc | 50 | 0.61 | 0.59 | 0.72 | 0.24 | 0.71 |
|  | cls | 50 | 0.63 | 0.61 | 0.69 | 0.24 | 0.72 |
|  | cn | 60 | 0.62 | 0.61 | 0.67 | 0.23 | 0.69 |
|  | ncn | 60 | 0.62 | 0.57 | 0.80 | 0.30 | 0.75 |
| Proxi | and | 60 | 0.64 | 0.61 | 0.76 | 0.30 | 0.75 |
|  | btw | 50 | 0.69 | 0.67 | 0.78 | 0.36 | 0.77 |
|  | cc | 60 | 0.62 | 0.61 | 0.68 | 0.24 | 0.70 |
|  | cls | 50 | 0.57 | 0.53 | 0.72 | 0.20 | 0.67 |
|  | cn | 40 | 0.65 | 0.62 | 0.77 | 0.31 | 0.78 |
|  | ncn | 60 | 0.67 | 0.65 | 0.75 | 0.33 | 0.77 |
| RMT | and | 50 | 0.66 | 0.64 | 0.75 | 0.32 | 0.78 |
|  | btw | 50 | 0.67 | 0.64 | 0.79 | 0.35 | 0.78 |
|  | cc | 20 | 0.68 | 0.66 | 0.78 | 0.35 | 0.78 |
|  | cls | 60 | 0.67 | 0.65 | 0.78 | 0.35 | 0.77 |
|  | cn | 60 | 0.68 | 0.66 | 0.75 | 0.33 | 0.77 |
|  | ncn | 60 | 0.68 | 0.66 | 0.76 | 0.34 | 0.78 |
| SparCC | and | 60 | 0.61 | 0.57 | 0.73 | 0.25 | 0.69 |
|  | btw | 60 | 0.68 | 0.66 | 0.75 | 0.34 | 0.75 |
|  | cc | 40 | 0.60 | 0.57 | 0.71 | 0.23 | 0.70 |
|  | cls | 60 | 0.66 | 0.65 | 0.72 | 0.29 | 0.73 |
|  | cn | 50 | 0.66 | 0.64 | 0.72 | 0.29 | 0.72 |
|  | ncn | 60 | 0.63 | 0.60 | 0.74 | 0.28 | 0.71 |

Second, we repeated the experiments described in the preceding paragraph but using CASS based on three node resilience measures as the Node Importance Scoring module in our NBBD framework. The performance of the resulting RF classifiers are reported in Table S6 and summarized in Table 3 for DS50. Table S6 shows that the highest AUC score of 0.79 can be reached using DS100 and graphs learned using Proxi (and 28 features) or SparCC (and 51 features) as well as using DS400 and graphs obtained using SparCC (and 54 features). Results in Table 3 suggest that the three CASS methods seem to need larger feature selection datasets in order to reach a predictive performance comparable to those obtained using traditional feature selection methods or NTPS methods. Unlike all other feature selection methods considered in this work, CASS methods do not require the user to provide the number of features to be selected from the input data as a parameter.

**Table 3.** Performance (highest AUC attained, and the smallest number of features chosen) by the top performing RF classifiers for combinations of different choices of Network Inference Method (NIM) and network-based feature selection using three resilience measures for Critical Attack Set Scoring (CASS). Results obtained using DS50 as the feature selection dataset.

| NIM | FSM | # Features | ACC | Sn | Sp | MCC | AUC |
| --- | --- | --- | --- | --- | --- | --- | --- |
| CoNet | CASS_I | 21 | 0.66 | 0.64 | 0.75 | 0.31 | 0.77 |
|  | CASS_T | 35 | 0.67 | 0.64 | 0.76 | 0.33 | 0.76 |
|  | CASS_V | 21 | 0.66 | 0.64 | 0.75 | 0.31 | 0.77 |
| MB | CASS_I | 6 | 0.51 | 0.47 | 0.68 | 0.12 | 0.61 |
|  | CASS_T | 33 | 0.57 | 0.53 | 0.73 | 0.21 | 0.65 |
|  | CASS_V | NA | NA | NA | NA | NA | NA |
| Proxi | CASS_I | 11 | 0.65 | 0.63 | 0.72 | 0.28 | 0.72 |
|  | CASS_T | 39 | 0.65 | 0.64 | 0.70 | 0.27 | 0.73 |
|  | CASS_V | 1 | 0.25 | 0.07 | 0.96 | 0.04 | 0.51 |
| RMT | CASS_I | 8 | 0.49 | 0.46 | 0.58 | 0.03 | 0.52 |
|  | CASS_T | 12 | 0.56 | 0.52 | 0.72 | 0.19 | 0.64 |
|  | CASS_V | 3 | 0.64 | 0.64 | 0.61 | 0.21 | 0.62 |
| SparCC | CASS_I | 117 | 0.66 | 0.64 | 0.74 | 0.31 | 0.76 |
|  | CASS_T | 125 | 0.66 | 0.64 | 0.72 | 0.30 | 0.76 |
|  | CASS_V | NA | NA | NA | NA | NA | NA |

In summary, our results suggest that the five NIMs, except MB^26^, can be successfully used in our NBBD framework for identifying discriminative features (i.e., potential IBD biomarkers) from metagenomics data. Our results also show that network-based feature selection methods are comparable to some commonly used traditional feature selection methods including the widely used RFFI. Moreover, with small size feature selection datasets, network-based feature selection methods applied to RMT graphs outperform traditional feature selection methods.

### Performance of Hybrid Feature Selection Methods

Preliminary results reported in an early version of this work (Fig. 4 in Abbas et al.^20^) show that only 12 OTUs were shared among the three subsets of 30 biomarkers determined using RFFI and two instances of the NBBD framework. Therefore, we hypothesize that the feature importance scores estimated using RFFI and the best instances of our NBBD framework are complementary with each other. To test this hypothesis, we developed a hybrid feature selection method that returns the product of RFFI and NBBD based on NTPSs as combined feature importance. Results for the hybrid method are reported for each of the five FSDS using graphs generated by five NIMs and instances of the NBBD framework using six topological properties for the NTPS approach and the top *k* ∈ {10, 20, 30, 40, 50, 60} biomarkers in Tables S7-S11 and the top performing RF classifiers using DS50 are reported in Table 4.

**Table 4.** Performance of the top performing RF classifiers (with the highest AUC and using the smallest number of features) for combinations of different choices of Network Inference Method (NIM) and hybrid feature selection based on RFFI and different properties for Node Topological Property Scoring. All results were obtained using DS50 as the feature selection dataset.

| GIM | FS Method | # Features | ACC | Sn | Sp | MCC | AUC |
| --- | --- | --- | --- | --- | --- | --- | --- |
| CoNet | and | 30 | 0.68 | 0.65 | 0.79 | 0.36 | 0.79 |
|  | btw | 20 | 0.65 | 0.61 | 0.79 | 0.33 | 0.78 |
|  | cc | 60 | 0.66 | 0.64 | 0.74 | 0.31 | 0.74 |
|  | cls | 40 | 0.62 | 0.59 | 0.77 | 0.29 | 0.74 |
|  | cn | 40 | 0.66 | 0.64 | 0.75 | 0.32 | 0.76 |
|  | ncn | 40 | 0.67 | 0.65 | 0.75 | 0.32 | 0.76 |
| MB | and | 20 | 0.73 | 0.72 | 0.76 | 0.40 | 0.82 |
|  | btw | 40 | 0.66 | 0.64 | 0.78 | 0.33 | 0.78 |
|  | cc | 20 | 0.66 | 0.62 | 0.79 | 0.34 | 0.77 |
|  | cls | 40 | 0.65 | 0.64 | 0.72 | 0.29 | 0.77 |
|  | cn | 10 | 0.69 | 0.68 | 0.74 | 0.34 | 0.76 |
|  | ncn | 20 | 0.65 | 0.61 | 0.80 | 0.34 | 0.79 |
| Proxi | and | 50 | 0.68 | 0.66 | 0.77 | 0.35 | 0.78 |
|  | btw | 30 | 0.69 | 0.66 | 0.82 | 0.39 | 0.79 |
|  | cc | 50 | 0.65 | 0.62 | 0.77 | 0.31 | 0.78 |
|  | cls | 50 | 0.67 | 0.63 | 0.83 | 0.37 | 0.79 |
|  | cn | 40 | 0.62 | 0.60 | 0.70 | 0.24 | 0.73 |
|  | ncn | 40 | 0.68 | 0.65 | 0.80 | 0.36 | 0.79 |
| RMT | and | 60 | 0.68 | 0.64 | 0.80 | 0.36 | 0.79 |
|  | btw | 40 | 0.64 | 0.60 | 0.80 | 0.32 | 0.78 |
|  | cc | 40 | 0.69 | 0.65 | 0.81 | 0.38 | 0.82 |
|  | cls | 50 | 0.69 | 0.66 | 0.80 | 0.37 | 0.80 |
|  | cn | 40 | 0.64 | 0.60 | 0.80 | 0.32 | 0.76 |
|  | ncn | 50 | 0.68 | 0.65 | 0.81 | 0.37 | 0.80 |
| SparCC | and | 30 | 0.67 | 0.64 | 0.78 | 0.34 | 0.80 |
|  | btw | 40 | 0.70 | 0.68 | 0.78 | 0.37 | 0.79 |
|  | cc | 30 | 0.66 | 0.63 | 0.79 | 0.34 | 0.78 |
|  | cls | 30 | 0.67 | 0.64 | 0.80 | 0.36 | 0.78 |
|  | cn | 50 | 0.67 | 0.63 | 0.82 | 0.36 | 0.80 |
|  | ncn | 40 | 0.66 | 0.62 | 0.81 | 0.34 | 0.79 |

Table S8 reports the results for RF classifier using hybrid feature selection based on instances of the NBBD framework applied to MB graphs and shows that the two best performing RF classifiers with AUC scores of 0.82 and 0.81 are obtained using the ‘and’ property and the top 10 and 20 features (respectively). Interestingly, these two classifiers were trained using features determined using MB graphs inferred from DS50. This is a substantial improvement in performance compared with the RF model trained using RFFI and features determined using DS50 (see Table 1) which has an AUC score of 0.80 using 50 features. In addition, several RF models with AUC scores higher than 0.80 were obtained using Proxi, RMT, and SparCC graphs (see Tables S7-S11).

Table 4 summarizes the results in Tables S7-S11 by reporting the top performing RF classifiers obtained using DS50 (i.e., the smallest feature selection dataset). In this table, two RF classifiers using MB and RMT graphs have equal AUC scores of 0.82. Several RF classifiers reached an AUC score of 0.80, but only the model based on SparCC graphs is using a small number of features. Comparing results in Tables 2 and 4 suggests that the RF classifiers using hybrid feature selection outperform counterpart RF classifiers using NTPS only in terms of predictive performance and/or number of features used to train the models.

### Analysis of Top Performing Models and the Identified IBD Biomarkers

Table 5 compares the performance of the top RF classifiers obtained using traditional feature selection and hybrid feature selection methods evaluated in our experiments. Using a hybrid scoring method combining RFFI (estimated from DS50) and ‘and’ scores (determined from MB graphs), a RF classifier trained using the top 20 features outperforms the best RF developed using RFFI (estimated from DS400) in four out of five performance metrics. Table 6 shows the AUC scores for these three models using different FSDSs. Since feature selection datasets are nested (i.e., *DS*50 ⊂ *DS*100 ⊂ *FSD*200 ⊂ *DS*300 ⊂ *DS*400), we expect feature selection methods to return the same or better subset of features as we increase the size of the FSDS used. Our expectation is almost realized using the RFFI method, except that there is a drop in AUC score when DS300 is used. On the other hand, our expectation is violated using the hybrid feature selection methods. The highest AUC score is observed using DS50, and increasing the size of the FSDS leads to a drop in classifier performance. This suggests that NIMs such as MB and RMT might be highly unstable to changes in the input data. In other words, networks constructed from DS50 and DS400 (as an example) are substantially different. For instance, Fig. S1 compares the four MB graphs generated using the MB method from IBD and healthy samples in DS50 and DS400. We found that MB constructs two networks (over the same set of nodes) but with a minimal overlap in edges from DS50 and DS400 data. In the absence of the ground truth, we can not determine which network is closer to reality. However, our results show that graphs inferred from DS50 allow our NBBD framework to identify a better set of features.

**Table 5.** Performance comparison of top three RF classifiers obtained using traditional feature selection and hybrid feature selection methods.

| NIM | FSDS | FS Method | # Features | ACC | Sn | Sp | MCC | AUC |
| --- | --- | --- | --- | --- | --- | --- | --- | --- |
| NA | DS400 | RFFI | 20 | 0.69 | 0.68 | 0.76 | 0.36 | 0.80 |
| MB | DS50 | RFFI × and | 20 | 0.73 | 0.72 | 0.76 | 0.40 | 0.82 |
| RMT | DS50 | RFFI × cc | 40 | 0.69 | 0.65 | 0.81 | 0.38 | 0.82 |

**Table 6.** AUC scores for top three RF classifiers obtained using RFFI feature selection and two hybrid feature selection methods, MB_and and RMT_cc, using different feature selection datasets.

| FSDS | RFFI | MB_and | RMT_cc |
| --- | --- | --- | --- |
| DS50 | 0.76 | 0.82 | 0.82 |
| DS100 | 0.78 | 0.76 | 0.79 |
| DS200 | 0.79 | 0.75 | 0.77 |
| DS300 | 0.77 | 0.79 | 0.77 |
| DS400 | 0.80 | 0.78 | 0.75 |

Fig. 2 shows the Venn diagram of unique and shared OTUs among the three subsets of features used for training the top three models in Table 5. We found that the number of unique OTUs in each subset is 7, 3, and 18 for RFFI, MB_and, and RMT_cc sets, respectively. Interestingly, 17 out of the 20 features in MB_and are also in RMT_cc and 8 out of these 17 common OTUs are also shared with RFFI. Table S12 lists the OTUs in these three sets of selected features. We further conducted downstream statistical analysis of the common 8 OTUs which are highlighted in bold in Table S12. More precisely, we assessed the significance of the difference between the medians of sample normalized relative abundance of these OTUs in IBD and healthy populations using the Kruskal-Wallis nonparametric test (Figures S2-S6). Analysis of DS400 (Fig. S6) shows significantly higher abundance of (Aggregatibacter, Fusobacterium, and Sutterella) in IBD samples relative to healthy samples. The increase of Aggregatibacter genus in IBD samples has been reported in several recent studies^44,45^. Also, the high abundance of Fusobacterium in IBD samples has been suggested as a biomarker in several studies^3,46^. Sutterella spp. have been frequently associated with several human diseases including autism and IBD^47,48^. However, other studies^49,50^ have suggested that Sutterella spp. are unlikely to play a role in the pathogenesis of IBD. Fig. S6 also shows significant decreases in Roseburia, Dialister, and Clostridiales. These three biomarkers have been repeatedly reported in previous studies^51^–53. Finally, results of our statistical analysis reported in Fig. S6 suggest that two of our top identified genera biomarkers, Bacteroides and Oscillospira, have no significant differences in IBD and control samples. Bacteroides is a dominant and biologically important bacteria genus in the microbiota of the human gastrointestinal tract^54^ and Oscillospira is an under-studied bacterial genus that is hard to cultivate but is consistently being identified in several human gut microbiota association studies^55^. This highlights the need for developing more sophisticated differential abundance tests that take into account the sparsity and compositional nature of metagenomics data.

**Figure 2.**
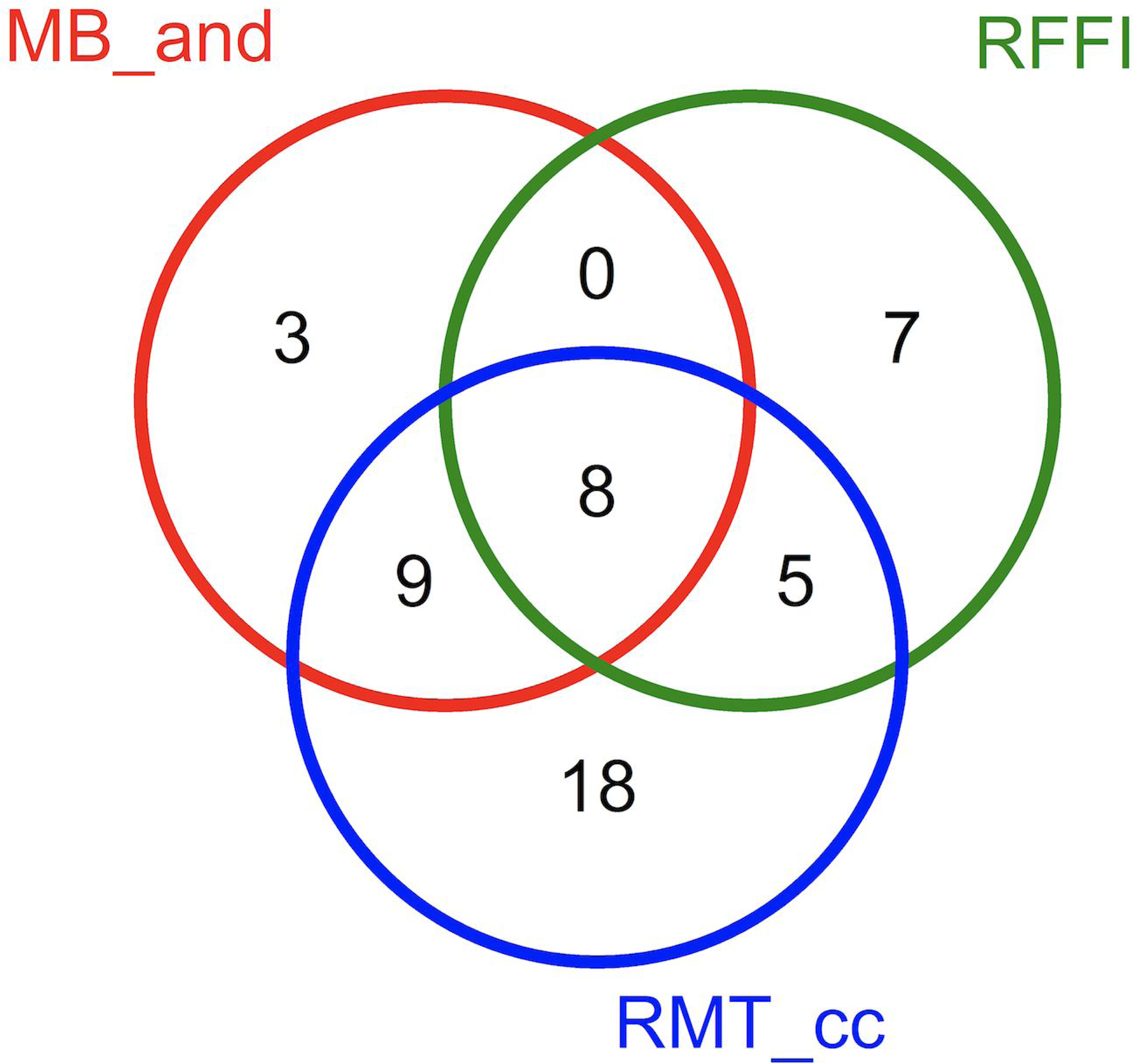
Venn diagram of unique and shared features selected using RF Feature Importance (RFFI), network-based feature selection applied to MB (RMT) networks and using ‘and’ (‘cc’) for node importance scoring.

Sensitivity analysis of Kruskal-Wallis and Mann-Whitney nonparametric tests against the number of samples analyzed has been conducted using all variants of FSDS. The complete results of this analysis is reported in supplementary figures S2-S11. Surprisingly, both tests failed to show any significant differences between IBD and healthy groups using DS50. Overall, the results from the two nonparametric tests are in agreement with each other, and our results suggest that at least 100 samples are needed for each group in order to demonstrate significant differences in the abundances of six out of the top eight identified biomarkers.

## Discussion

The past decade has witnessed a revolution in microbiology and microbiome research. Advances in sequencing technologies and computational techniques coupled with large scale collaborative efforts such as Human Microbiome Project (HMP)^56^ and American Gut Project^57^ have generated unprecedented amounts of metagenomics data. Analysis and interpretation of such data presents many statistical and computational challenges^58,59^. One such challenge has to do with the reliable identification of biomarkers (in the form of species, genes, or pathways) that differentiate between two or more phenotypes^12^.

To address this challenge, we have developed NBBD, a novel metagenomics system biology framework for microbial biomarker discovery. The NBBD framework integrates network analysis and machine learning approaches for reliable identification of biomarkers from metagenomics data. Given two OTU tables corresponding to two phenotypes, NBBD uses any existing tool for constructing phenotype specific networks from the data. Depending on the tool used, these networks model the interactions, the correlations, or the proximity relationships between microbes. Next, the nodes are scored using different scoring methods that quantify the extent to which the nodes contribute to differences in the topological properties of the nodes in the two networks. The *k* top-scoring nodes are used as the set of selected features to train and test classifier using machine learning. We conducted extensive experiments to evaluate the NBBD framework, configured using five different network inference tools and nine different node importance scoring methods, using a large dataset from a cohort of 657 IBD and 316 healthy healthy pediatric metagenomics biopsy samples, respectively.

Although several tools for constructing microbial ecology networks from metagenomics data have been developed, they leave considerable room for improvement^12,60^. For example, Weiss et al.^60^ benchmarked the performance of eight correlation detection strategies on simulated and real metagenomics data and showed significant inconsistency (in terms of number of edges) among graphs inferred using different tools. Using simulated data, they showed that all of the tools exhibited extremely low precision (below 0.20). That is, for every identified true edge, there are at least four false positive edges in the constructed network. While the five network construction tools considered in our study are among the top performing tools in Weiss et al.^60^, they are far from perfect. It is indeed remarkable that the noisy networks produced by such tools can be used to reliably identify discriminative features and to identify potential IBD biomarkers.

In this study, we performed experiments to examine the sensitivity of classifiers to the number of samples in the feature selection dataset. To facilitate fair comparison between classifiers, we used the entire training data for training the classifiers using the features determined based on different subsets of the training data. Our results suggest that traditional feature selection methods fail to determine a minimal subset of discriminative features from small feature selection datas ets. Interestingly, we found that several network-based feature selection methods returned a minimal subset of discriminative features using the smallest feature selection dataset, DS50. These findings highlight one of the reasons network-based feature selection should be used. Mainly, by mapping the feature selection data into graphs, we overcome several challenges in the input data, including a small number of samples, sparsity, and high-dimensionality. Another reason for using network-based feature selection is that it opens up the possibility of developing a variety of novel feature selection methods based on a broad and rich collection of well-developed graph mining algorithms. For example, in this work, we showed how to develop network-based feature selection methods using virtually any vertex topological property and also using a graph clustering algorithm (NBR-Clust). In particular, the CASS method (derived from NBR-Clust) determines the optimal number of features seamlessly. More methods could be developed using vertex similarity algorithm (e.g., SimRank^61^ and ASCOS^62^), graph similarity algorithms (e.g., DeltaCon^63^), and network-based anomaly detection methods^64^. Our ongoing work aims to explore the utility of these algorithms for developing more sophisticated Node Importance Scoring (NIS) modules for the NBBD framework.

Our sensitivity analysis also revealed that the microbial ecology networks constructed using state-of-the-art network construction methods are highly sensitive to the data samples used to construct the network. Needless to say, this lack of stability of network construction algorithms has serious implications for subsequent biological interpretation of microbial ecology networks, and in the contest of our work, the reliability of the biomarkers discovered from analysis of microbial ecology networks. In order for the predictive models trained using the features selected using network-based feature selection methods) to be reliable, we need to ensure the feature selection methods have a high degree of stability with respect to changes in the underlying network. Note that the stability of feature selection algorithms is a function of both the properties of the algorithm itself as well as the data supplied to the algorithm. Hence, improvements are needed on both fronts.

Fundamentally, constructing microbial ecology networks from metagenomic data requires determining the correlation or similarity between (abundances of) microbial taxa from a relatively small number of metagenomic samples. This problem is not fundamentally different from the problem of determining gene co-expression networks from gene expression data^65^, or that of determining functional brain networks from fMRI data^66^. All of these applications present some shared challenges: In most cases, the number of features (genes, brain regions, microbial taxa) far exceed the number of data samples; It is generally impossible, without making additional assumptions or incorporating domain knowledge, to distinguish between direct and indirect correlations; The choice of the correlation or similarity measure is often application-dependent. Methods for microbial ecology network estimation from metagenomic data could benefit greatly from recent advances in high dimensional correlation matrix estimation^67–70^. Work in progress is aimed at evaluating the applicability of such methods in constructing stable microbial ecology networks from metagenomic data.

## Conclusions

We have proposed a novel Network-Based Biomarker Discovery (NBBD) framework for detecting disease biomarkers from metagenomics data. NBBD consists of two major customizable modules: A network inference module, for constructing microbial ecology networks from OTU tables extracted from the metagenomic data for the phenotypes of interest; and a node importance scoring module, which compares the resulting phenotype-specific networks and scores the nodes based on different measures of the node’s contribution to the differences between the networks.

We have evaluated the proposed NBBD framework, using five different network construction methods, in combination with nine different node importance scoring methods, on a large dataset from a cohort of 657 IBD and 316 healhy pediatric metagenomics biopsy samples. Our results show that NBBD, when used to train predictive models for IBD diagnosis from metagenomic data, is very competitive with some of the state-of-the-art feature selection methods including the widely used method based on random forest feature importance scores. Our results further show that a hybrid approach that combines NBBD scores and the random forest feature importance scores yields further improvements in performance. Furthermore, the proposed method is able to achieve its best observed performance using only only 50 samples for feature selection. Work in progress is aimed at further improving the two key components of NBBD, e.g., by incorporating recent advances in high dimensional correlation matrix estimation^67–70^ to improve the reliability and the stability of the resulting networks, exploring improved node scoring methods. Other promising directions for future research include systematic evaluation of the NBBD framework for biomarker discovery from different types of omics data, integrative analyses of multi-omics data^71,72^, e.g., using information-preserving low-dimensional network embeddings^73^.

## Acknowledgements

V.H. was supported in part by the National Center for Advancing Translational Sciences, National Institutes of Health, through the Grant UL1 TR000127 and TR002014 in support of the Penn State Clinical and Translational Scinces Institute, by the National Science Foundation, through the grants 1518732, 1640834, and 1636795; the Penn State Center for Big Data Analytics and Discovery Informatics, the Penn State Institute for Cyberscience, the Edward Frymoyer Endowed Professorship in Information Sciences and Technology at Penn State, and the Pratiksha Trust, through the Sudha Murty Distinguished Visiting Chair in Neurocomputing and Data Sciences at the Indian Institute of Science. Y.E. was supported in part by the Center for Big Data Analytics and Discovery Informatics at the Pennsylvania State University and the Penn State Clinical and Translational Sciences Institute. The work of T.L was supported in part by the institutional match on the Predoctoral Training Grant T32-LM012415 in Biomedical Data Sciences from the National Library of Medicine, National Institutes of Health. The publication costs were covered by Qatar Computing Research Institute. The content is solely the responsibility of the authors and does not necessarily represent the official views of the funding agencies.

## Author contributions statement

T.O. and Y.E. conceptualized the study and designed the experiments, M.A. J.M. T.L. Y.E. conducted the experiment(s), H.B. T.O. V.H. Y.E. analyzed and interpreted the results. M.A. J.M. T.O. Y.E. drafted the manuscript. V.H. and Y.E. edited the manuscript. All authors read and approved the final version of the manuscript.

