## Supplementary material for "Biomarker discovery in inflammatory bowel diseases using network-based feature selection": Supplementary_Figures_S1-S11.docx

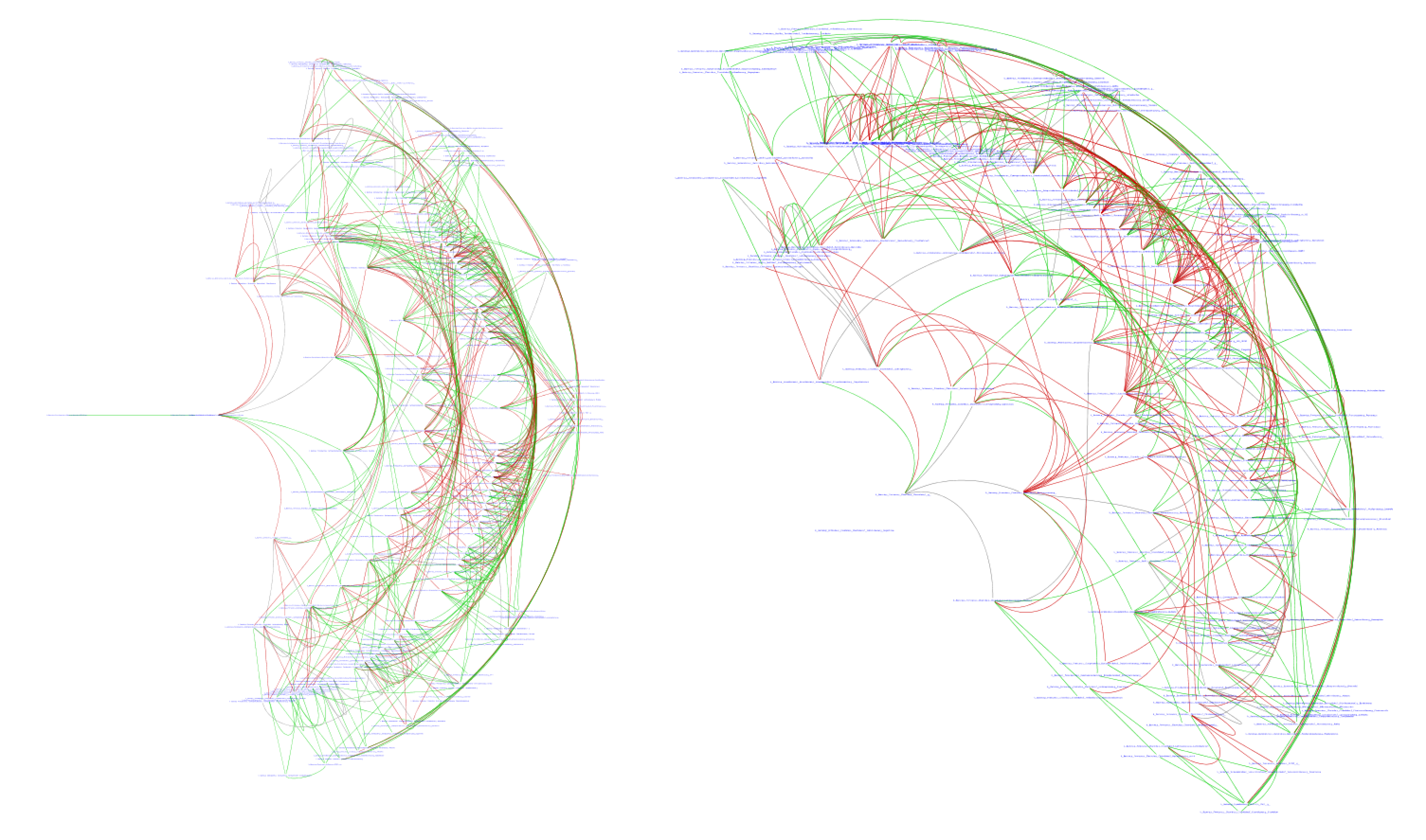

**Supplementary Figure S1: Comparisons between IBD (left) and healthy (right) networks inferred using MB tool applied to FSDS50 (green) and FSDS400 (red). Gray edges represent common edges between the two compared networks.**

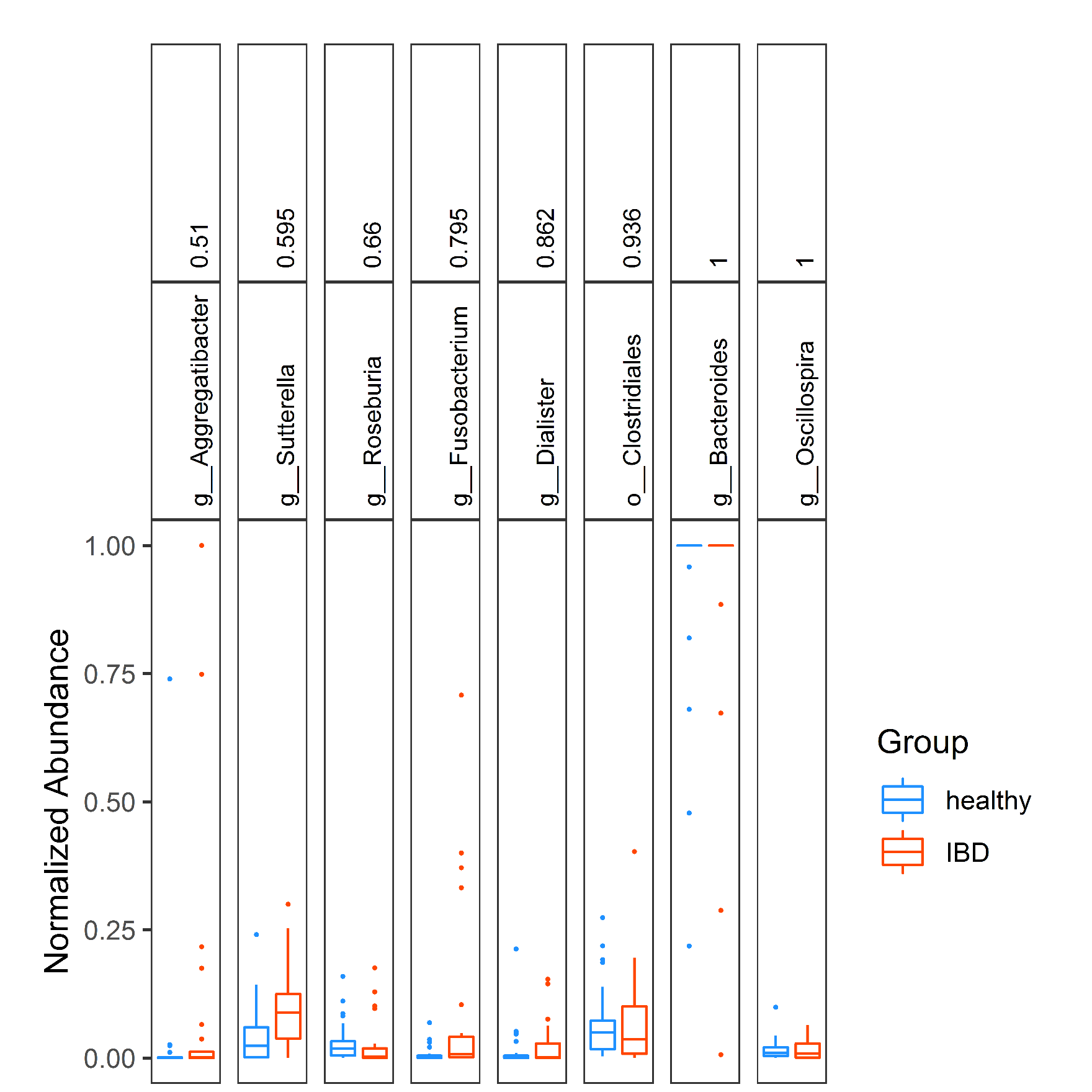

**Supplementary Figure S2: Box plots of normalized abundance for the eight common IBD biomarkers including p-values obtained using Kruskal-Wallis test of medians applied to FSDS50**

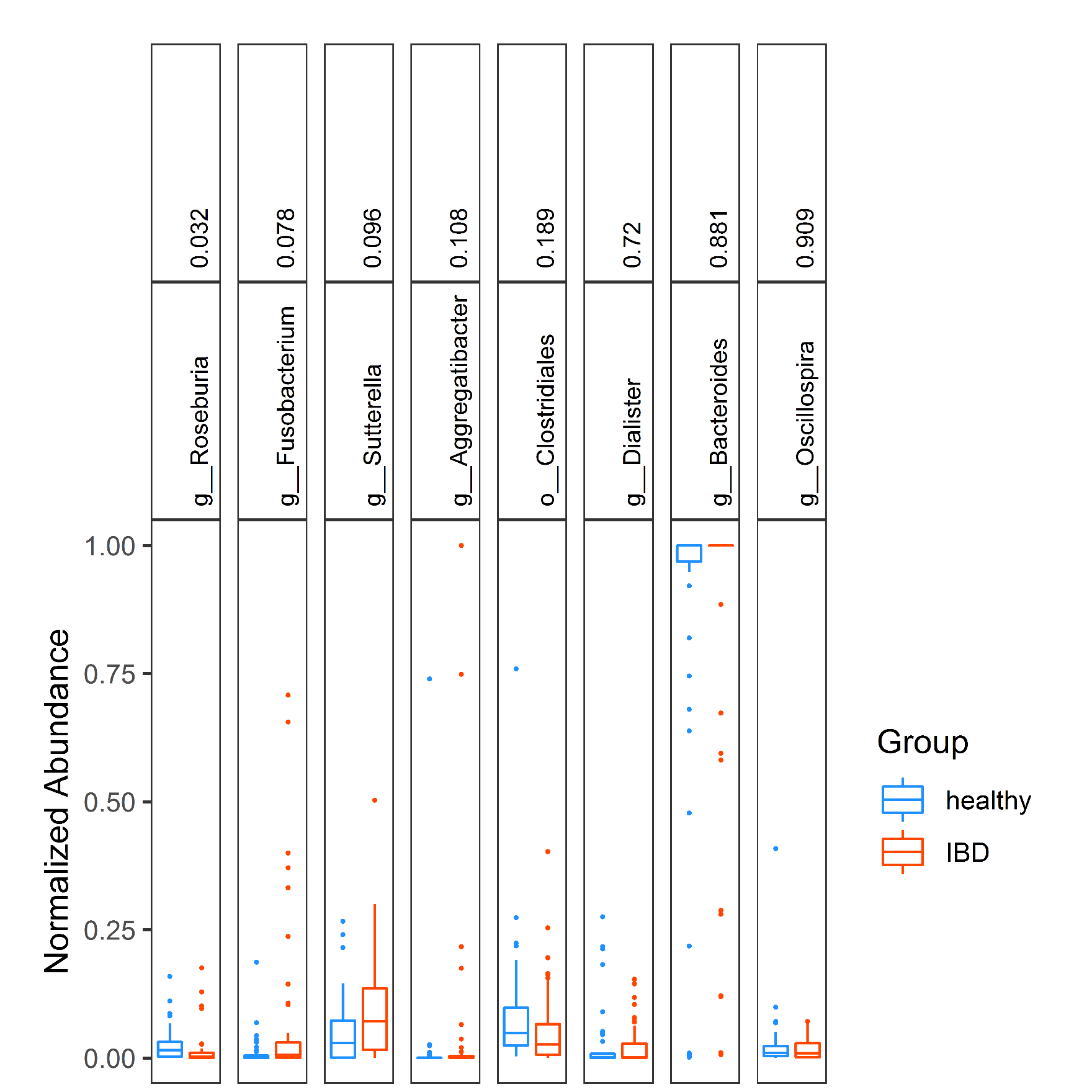

**Supplementary Figure S3: Box plots of normalized abundance for the eight common IBD biomarkers including p-values obtained using Kruskal-Wallis test of medians applied to FSDS100**

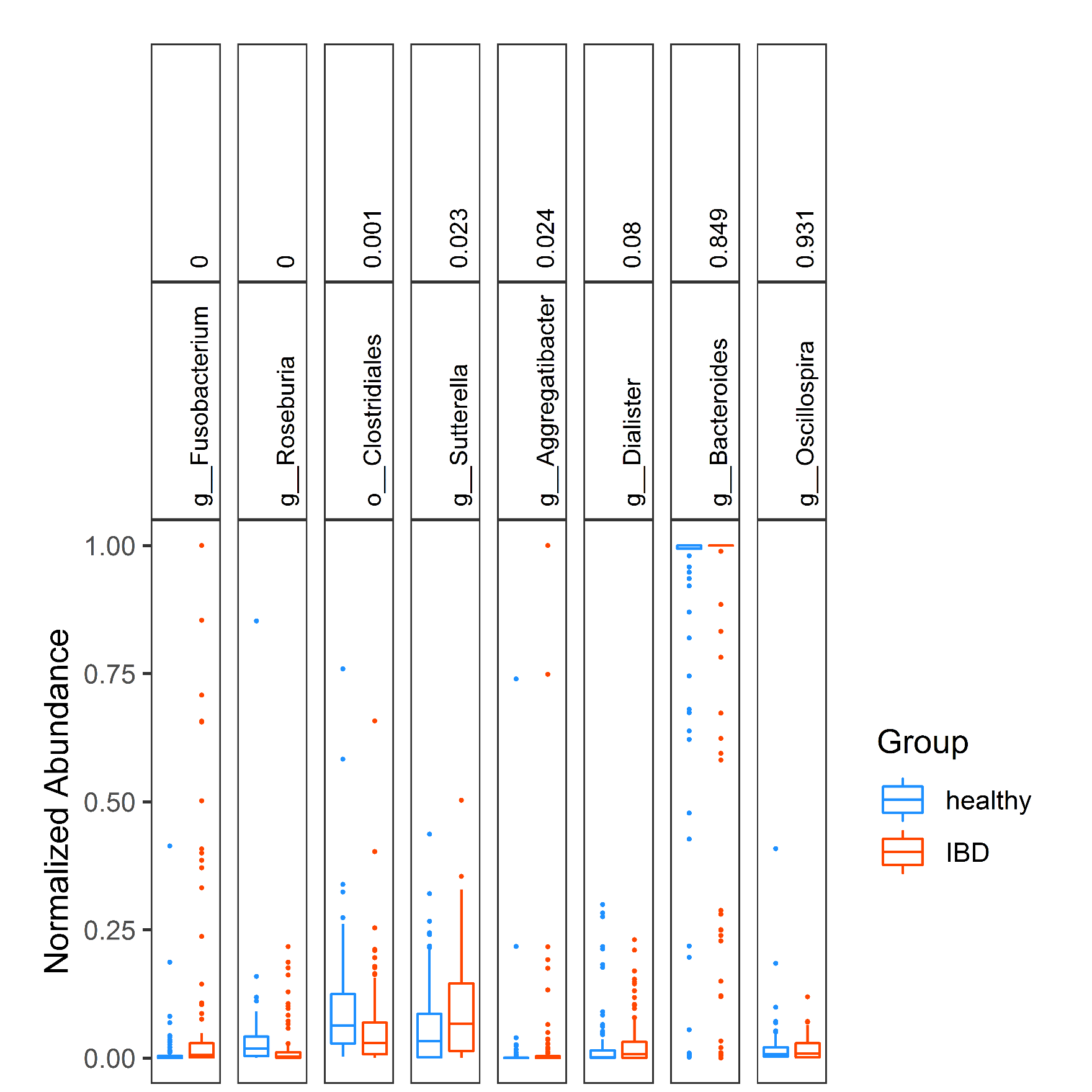

**Supplementary Figure S4: Box plots of normalized abundance for the eight common IBD biomarkers including p-values obtained using Kruskal-Wallis test of medians applied to FSDS200**

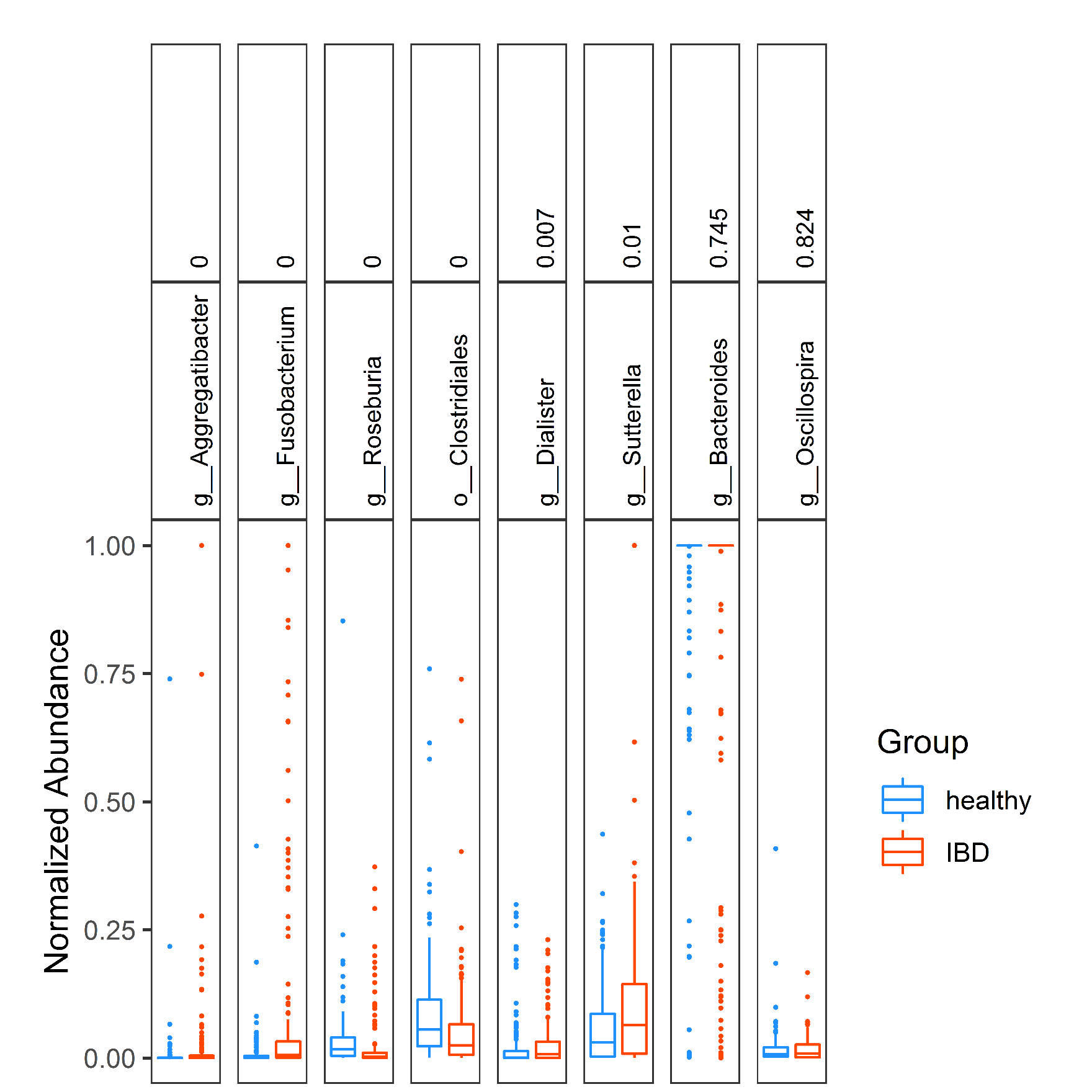

**Supplementary Figure S5: Box plots of normalized abundance for the eight common IBD biomarkers including p-values obtained using Kruskal-Wallis test of medians applied to FSDS300**

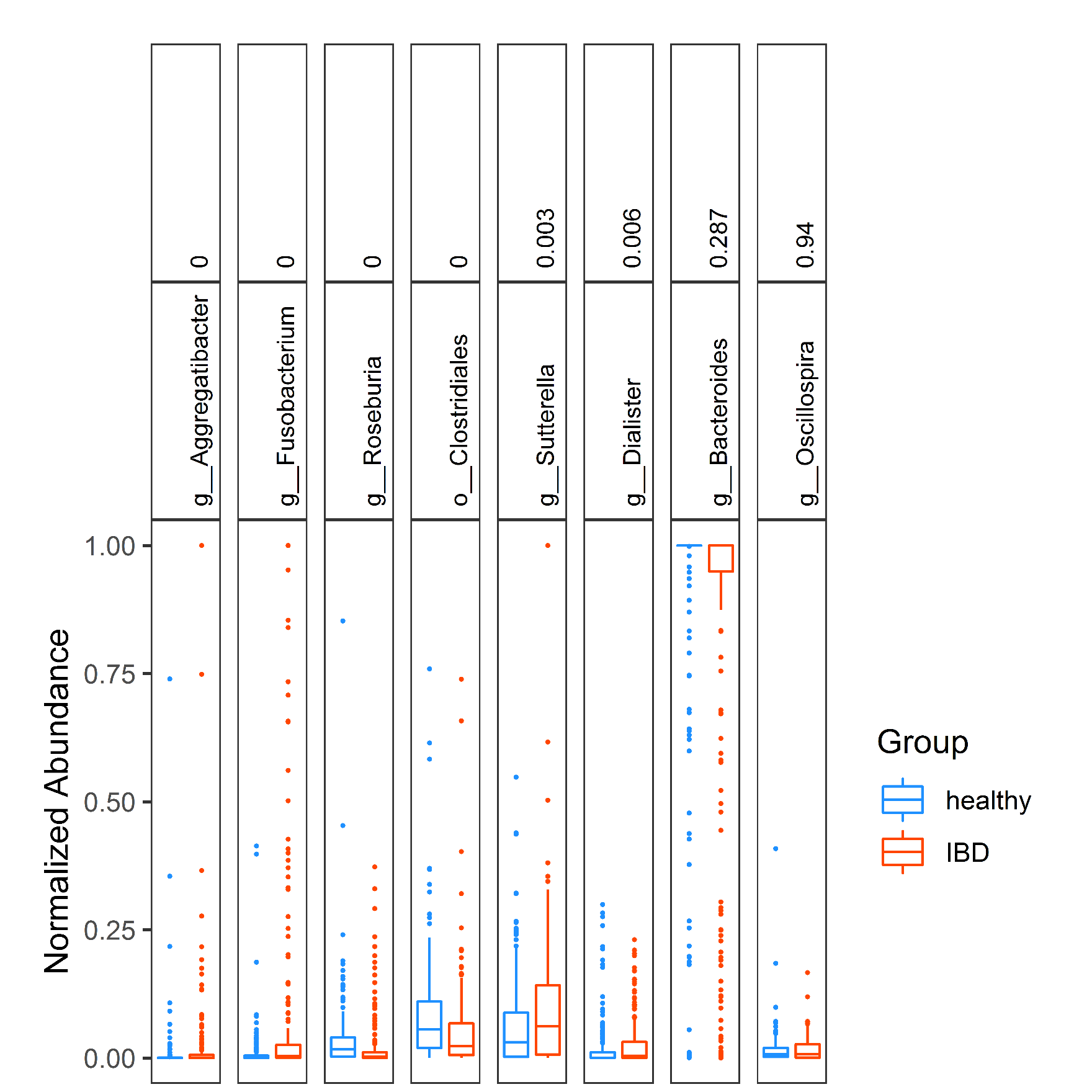

**Supplementary Figure S6: Box plots of normalized abundance for the eight common IBD biomarkers including p-values obtained using Kruskal-Wallis test of medians applied to FSDS400**

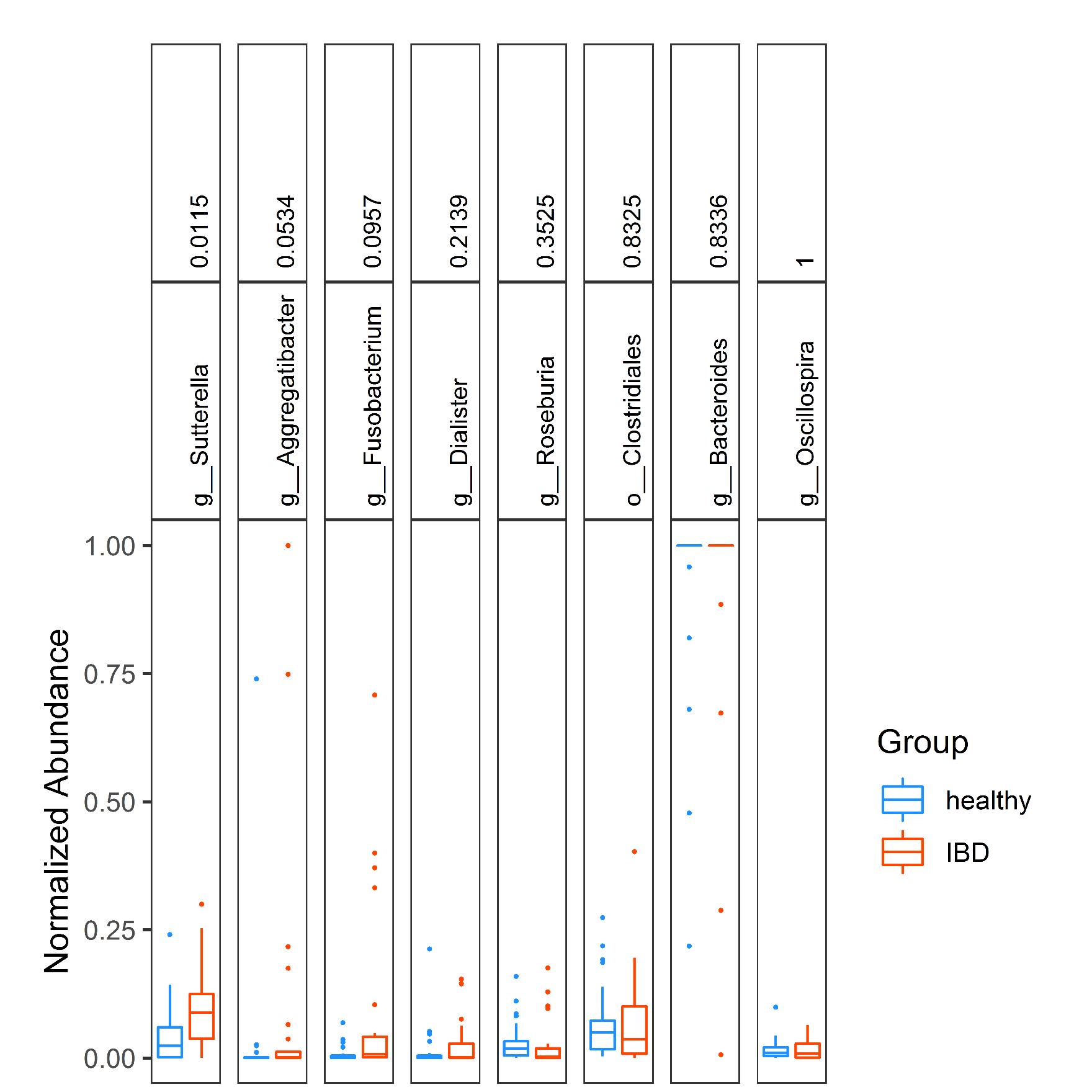

**Supplementary Figure S7: Box plots of normalized abundance for the eight common IBD biomarkers including p-values obtained using Mann-Whitney test of medians applied to FSDS50**

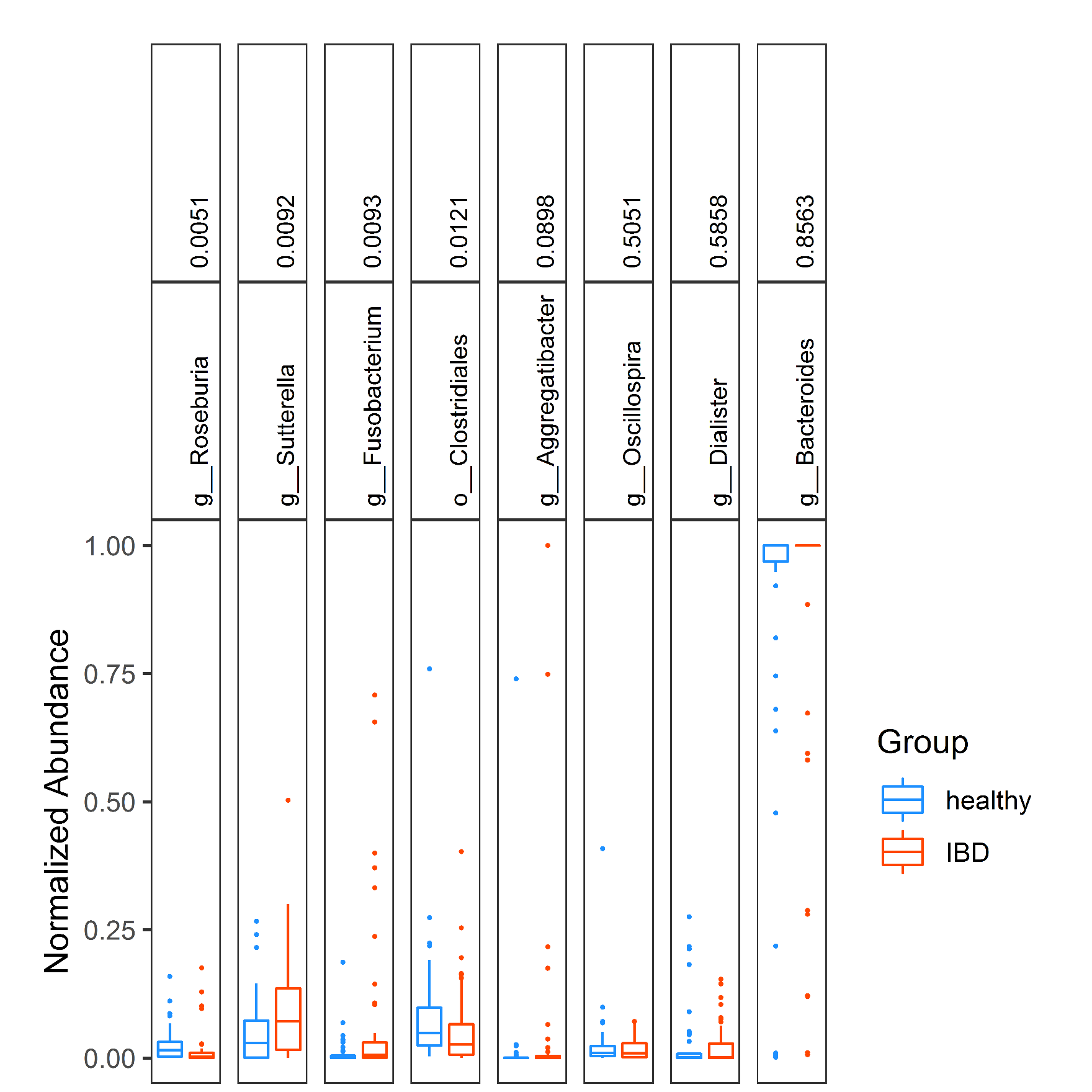

**Supplementary Figure S8: Box plots of normalized abundance for the eight common IBD biomarkers including p-values obtained using Mann-Whitney test of medians applied to FSDS100**

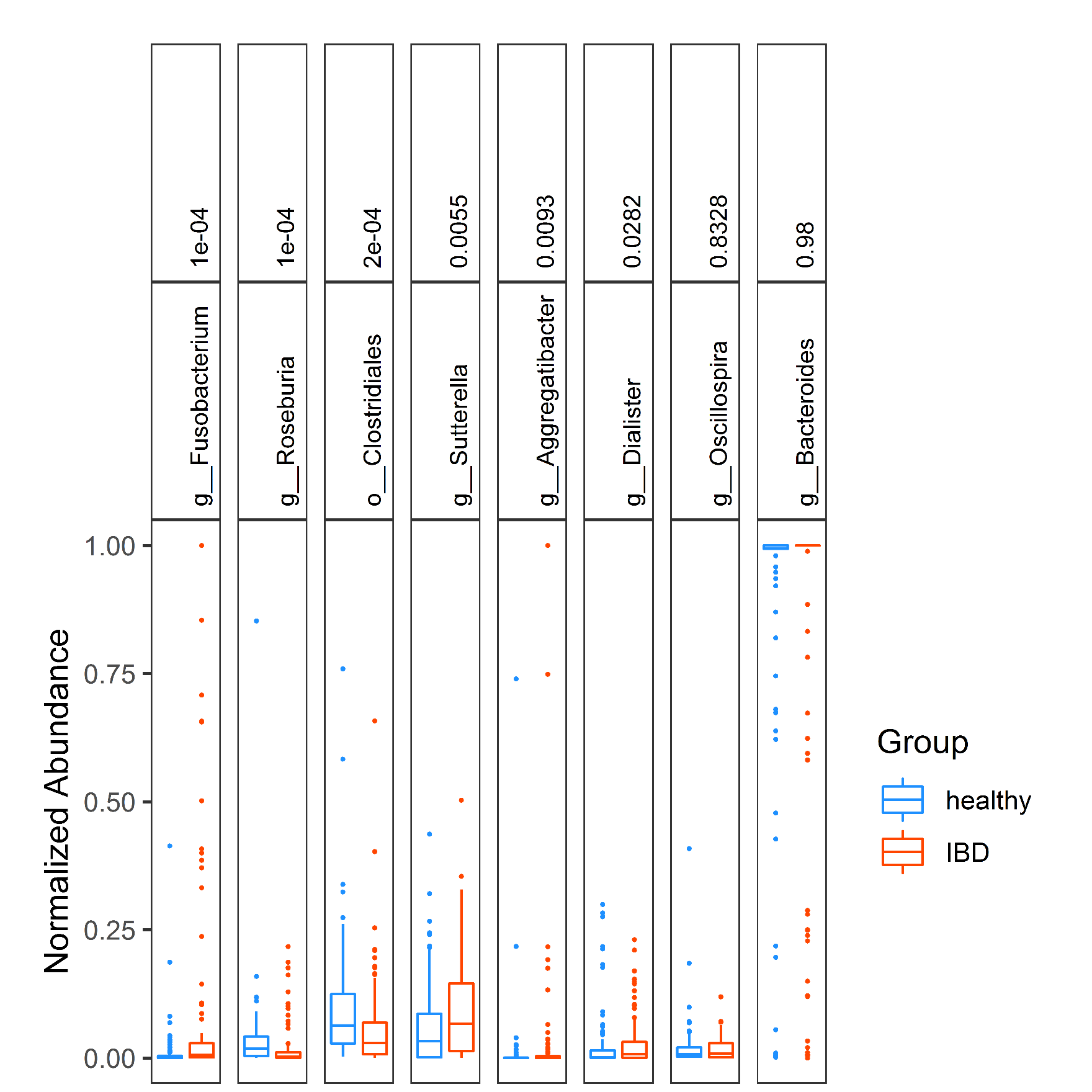

**Supplementary Figure S9: Box plots of normalized abundance for the eight common IBD biomarkers including p-values obtained using Mann-Whitney test of medians applied to FSDS200**

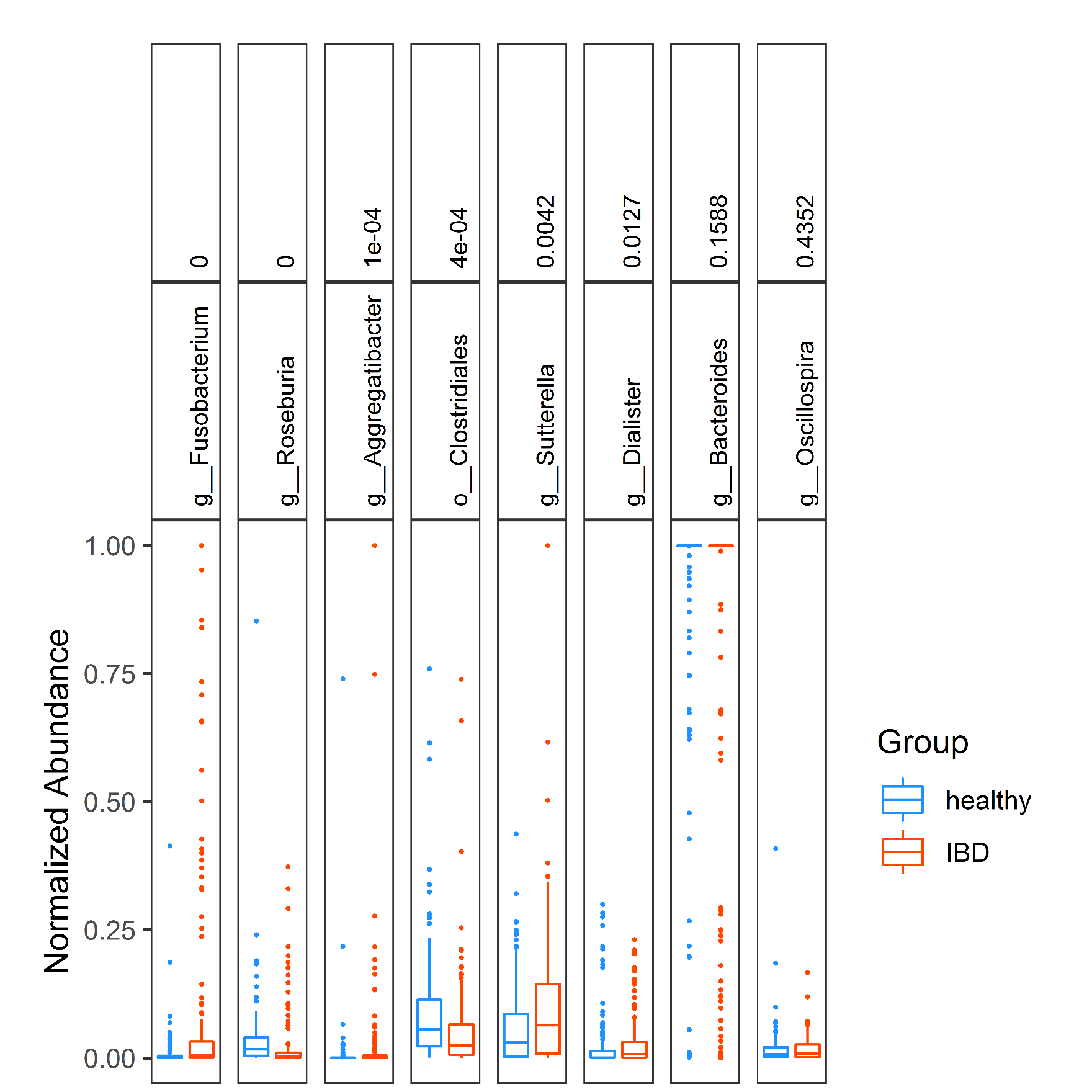

**Supplementary Figure S10: Box plots of normalized abundance for the eight common IBD biomarkers including p-values obtained using Mann-Whitney test of medians applied to FSDS300**

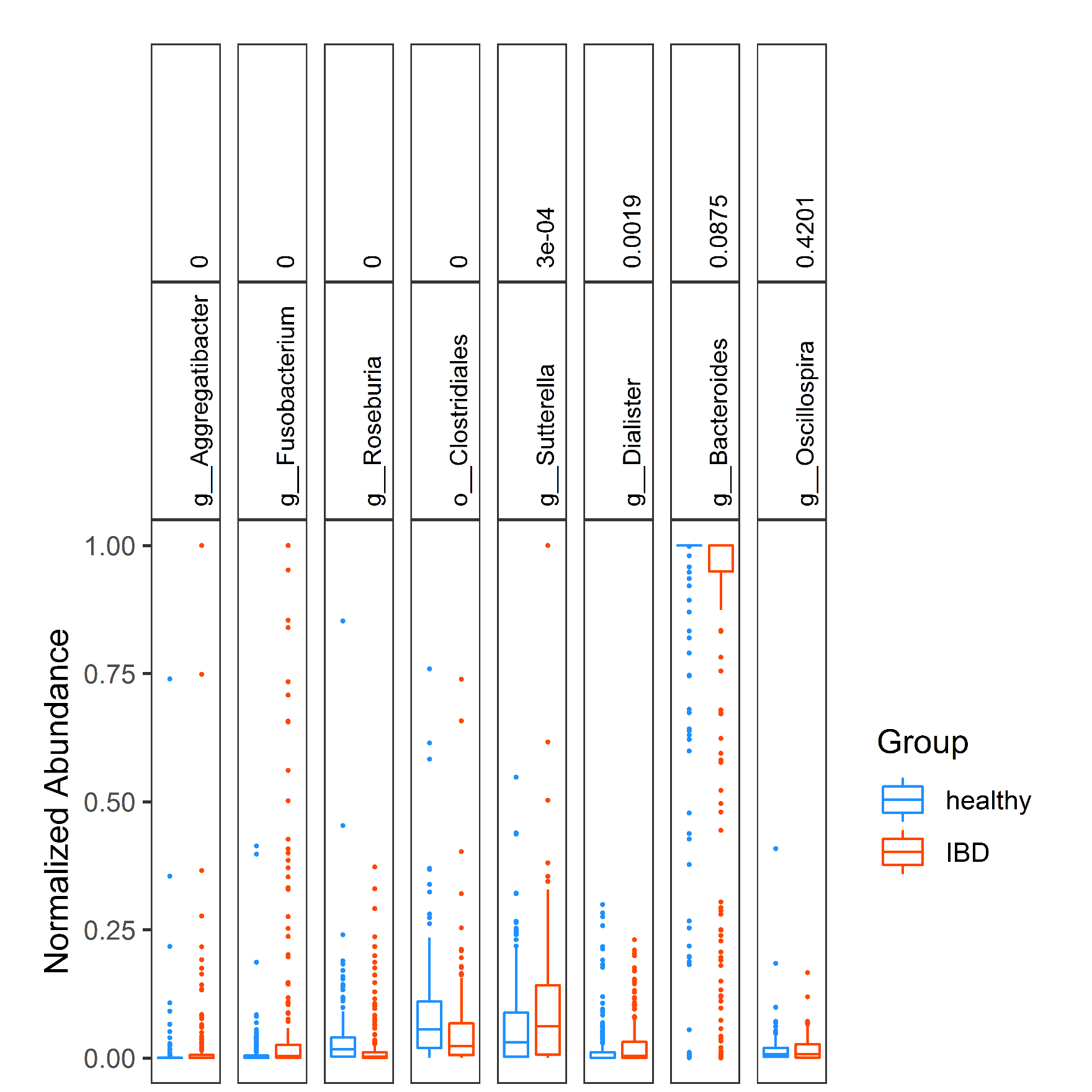

**Supplementary Figure S11: Box plots of normalized abundance for the eight common IBD biomarkers including p-values obtained using Mann-Whitney test of medians applied to FSDS400**
